# Population genomic scans reveal novel genes underlie convergent flowering time evolution in the introduced range of *Arabidopsis thaliana*

**DOI:** 10.1101/023788

**Authors:** Billie A. Gould, John R. Stinchcombe

## Abstract

A long-standing question in evolutionary biology is whether the evolution of convergent phenotypes results from selection on the same heritable genetic components. Using whole genome sequencing and genome scans, we tested whether the evolution of parallel longitudinal flowering time clines in the native and introduced ranges of *Arabidopsis thaliana* has a similar genetic basis. We found that common variants of large effect on flowering time in the native range do not appear to have been under recent strong selection in the introduced range. Genes in regions of the genome that are under selection for flowering time are also not enriched for functions related to development or environmental sensing. We instead identified a set of 53 new candidate genes putatively linked to the evolution of flowering time in the species introduced range. A high degree of conditional neutrality of flowering time variants between the native and introduced range may preclude parallel evolution at the level of genes. Overall, neither gene pleiotropy nor available standing genetic variation appears to have restricted the evolution of flowering time in the introduced range to high frequency variants from the native range or to known flowering time pathway genes.

## Introduction

A long-standing question in evolutionary biology is whether the evolution of convergent phenotypes results from selection on the same heritable genetic components. Are there multiple genetic mechanisms to produce adaptive phenotypes? Or, do correlations between traits and the pleiotropic constraint imposed by some genes and complex pathways confine evolution to using the same genetic mechanisms or loci? Answering these questions provides direct evidence of the role genetic constraint or flexibility in evolutionary responses. A fundamental goal in evolutionary biology is to understand which genes are subject to selection and how often, and to evaluate the general predictability of the evolutionary process at the phenotypic and genetic levels (Wood *et al*. 2005; Arendt and Reznick 2008; Elmer and Meyer 2011; Lee *et al*. 2014). Here, we show how parallel evolution in a key life history trait, flowering time, in the introduced range of the mouse-ear cress (*Arabidopsis thaliana*) utilizes novel loci, suggesting that numerous independent genetic mechanisms can be used to produce adaptive phenotypes.

Convergent evolution of locally adaptive traits can play a significant role in the successful establishment of plants outside their native ranges (Lee 2002; Bossdorf *et al*. 2005; Buswell *et al*. 2011). The existence of parallel geographic clines in the native and introduced ranges is often an indicator that similar climate-based selective pressures have promoted convergent local adaptation in both ranges. For example, North American populations of St. John’s Wort (*Hypericum perforatum*) exhibit clines in physiological traits, biomass, and fecundity that parallel patterns found in its native European range (Maron *et al*. 2004, 2007). Similarly, the invasive purple loosetrife has rapidly evolved geographic differences in time to flowering similar to those found in Europe (Montague *et al*. 2008; Colautti and Barrett 2013). As natural replicates of the evolutionary process, convergently evolved, native and introduced populations are often used as evidence of natural selection shaping phenotypic trends. They also provide a valuable scenario for testing to what extent selection acts repeatedly on the same traits, genes, and mutations. Theory (Orr 2005) and intuition suggest that parallel evolution at the genetic level will be common when there are relatively few possible genetic mechanisms to produce adaptive phenotypes, and rare when there are many genetic routes to adaptation. Adaptation from standing genetic variation in the introduced range is also predicted to occur first through selection on variants that are already high in frequency, and are (or were) adaptive (Barrett and Schluter 2008). To date, however, there have been few studies where the genetic basis of variation in convergently evolved traits has been compared between native and introduced populations.

For plants, flowering when environmental conditions are optimal is one of the most important traits for maximizing fitness, and much of what we know about the genetic pathways controlling flowering time has resulted from molecular work on laboratory strains of *Arabidopsis* (reviewed (Simpson and Dean 2002)). More recently, studies of naturally occurring *Arabidopsis* accessions have revealed a suite of both rare and common genetic variants that influence flowering time variation in the species native European range. Some mutations are common and control a large portion of the range-wide variation in phenotype while others are regionally restricted in their distribution. Null mutations affecting the interacting loci *Frigida (Fri) and Flowering Locus C* (FLC) explain 50-70% of flowering time variation under over-wintering conditions in some studies (Caicedo *et al*. 2004; Shindo *et al*. 2005). Common, high-impact variants also occur in the light sensing protein *Phytochrome C* (PHYC) (Balasubramanian *et al*. 2006; Samis *et al*. 2008) and in the transcription factor MADs affecting flowering 2 (MAF2) (Caicedo *et al*. 2009; Rosloski *et al*. 2010). Rare variants in FRI, FLC and other loci also contribute to differences between specific lines (Gazzani *et al*. 2003; Shindo *et al*. 2005; Li *et al*. 2014). Multiple-cross QTL experiments have yielded similar results as to the number of genomic regions that control flowering time variation between divergent European ecotypes. In a study of 13 F2 crosses between mainly European parental genotypes, Brachi *et al*. (Brachi *et al*. 2010) found 5 genomic regions containing upwards of 60 QTL implicated in flowering time variation, many of which were unique to a single cross. In a study of 17 F2 populations, Salomé *et al*. (SalomÉ *et al*. 2011) found flowering time variation was also linked to as few as 5 genomic regions. A recent genome-wide association study that included analysis of 23 separate flowering time related phenotypes identified significant SNPs in 7 loci, some of which affected known flowering time pathway genes (including *FRI* and *FLC*) and some of which did not (Aranzana *et al*. 2005; Atwell *et al*. 2010; Horton *et al*. 2012).

Mutations affecting flowering time variation in A. thaliana have been largely generated from studies that included few or no genotypes from the species introduced range in North America (but see (Atwell *et al*. 2010)). Only one study of which we are aware has directly compared flowering time differences between genotypes from the native and introduced ranges. Samis *et al*. (Samis *et al*. 2012) found similar longitudinal flowering time clines in lines from both Europe and North America. Under natural over-wintering conditions in a common garden, plants from more coastal populations flowered later in both Europe and North America. The cline was significant on both continents, but stronger in Europe (-0.20 days/degree longitude) than in North America (-0.09 days/degree longitude, Fig S1). In the introduced range, the cline is strongly correlated with total annual precipitation, and in these lines selection favors early flowering under low water conditions (Stock *et al*. 2015). As an initial test for a parallel genetic basis underlying flowering time clines in the native and introduced range, Samis and colleagues genotyped populations for 8 common polymorphisms across *Fri*, *FLC*, and *PHYC*. Some common alleles were present in the introduced range but none were strongly associated with flowering time phenotype. It remains unknown whether other less common flowering time alleles, novel mutations affecting flowering time pathway genes, or variants in other parts of the genome have been co-opted during local adaptation in the introduced range.

In the present study, we used a whole genome sequencing approach to search for evidence of selection associated with flowering time differences in introduced *Arabidopsis* genotypes. First, we identified SNPs genome-wide that show high levels of genetic differentiation between early and late flowering individuals, while controlling for population structure. We paired this with multiple diversity-based scans to identify potential regions of recent selection associated with flowering time variation. We then compared the results of these analyses to the functional categorization of the implicated regions and known flowering time genes and mutations from the native range. Collectively, our results suggest that neither pleiotropic constraint nor available standing genetic variation has constrained the evolution of flowering time in the species introduced range to selection on high frequency variants from the native range or to known flowering time pathway genes.

## Methods

### Plant material

Thirty-four North American *Arabidopsis* lines were selected for sequencing from among natural accessions previously phenotyped by Samis *et al*. (2012). The lines were obtained from the *Arabidopsis* Biological Resource Center (ABRC) and from recent collections from natural populations (Table S2). The lines were phenotyped in a rooftop common garden at the University of Toronto after overwintering under natural conditions (see (Samis *et al*. 2012)). We chose 17 late flowering and 17 early flowering lines for whole genome sequencing, using previous SNP data to choose lines with the maximum level of genetic diversity. The average flowering time difference between the two phenotypic groups was 6.7 days. In total the lines contain between 1 and 5 individuals from each of 14 different locations in the eastern United States. To generate plant material for sequencing, seeds were stratified in 0.15% agarose at 4 deg C for 3 days, and then germinated in soil in a growth chamber at 20 deg C under 16 hour days. DNA was extracted from young leaves using the Qiagen DNeasy Plant Mini kit. One barcoded sequencing library was constructed per individual using the Illumina TruSeq DNA sample prep kit. Library preparation and sequencing were performed at the McGill University and Genome Quebec Innovation Center (Montréal, Québec).

### Sequencing and variant detection

Whole genome sequencing was performed on two lanes of the Illumina HiSeq2000 platform using 100 bp paired end reads. We obtained an average of 24.4 million high quality reads per individual. We aligned reads back to the TAIR10 reference genome using Stampy (Lunter and Goodson 2011) with a predicted substitution rate of 0.007 (Mitchell-Olds and Schmitt 2006). On average 92.5% of reads could be mapped back to the reference per individual, generating an average of 14.6X coverage across the autosomes. We removed PCR duplicates from each alignment using Picard Tools v1.98 (“http://broadinstitute.github.io/picard” 2015) and used the Genome Analysis Toolkit (GATK, v2.7-2) for downstream filtering and variant calling. We identified insertions and deletions using GATK RealignerTargetCreator, and realigned reads around them using IndelRealigner. We created reduced alignment files using ReduceReads and called variants using Haplotype Caller. Variants were called using the set of all alignments in discovery mode with a strand call confidence of 30, and an emit confidence of 10. We filtered the initial set of called variants following guidelines provided by GATK best practices documentation. SNPs were excluded that had quality divided by read depth (QD) < 2.0, Fisher Strand Bias (FS) > 60.0, and mapping quality (MQ) < 40.0. Indels were removed that had QD < 2.0, FS > 200.0, and MQ < 40. Filtering removed about 6.7% of the original variants, leaving 2,453,132 autosomal variants in the data set. Downstream analyses were conducted using VCFtools (0.1.11), Python (v.2.7.5), and R (CORE Team 2012).

We compared the variants in our lines with results from two previous resequencing studies. We identified all variant sites with non-missing allele calls in at least half of the lines (n= 2,347,399 sites). These were compared with filtered variant calls published from the 1001 genomes project (Cao *et al*. 2011) http://1001genomes.org/data/MPI/MPICao2010/releases/2012_03_13/strains/ and variants from 19 genomes that make up the MAGIC nested association panel (Gan *et al*. 2011) (downloaded from http://mus.well.ox.ac.uk/19genomes/variants.tables/)

We validated the filtered set of variants against genotype calls at 135 snps previously genotyped in the same lines using Sanger sequencing (Samis *et al*. 2012). Two of the lines had a low validation rate (<60% of markers had matching calls) and we excluded these from further analysis (both were early flowering genotypes). Among the remaining 32 lines, concordance at homozygous sites between genotype calls using the two methods was 96.4%.

From the initial data set, we isolated a set of high confidence variants. We chose to retain only variants on the autosomes with no missing data, a minor allele frequency above 0.0625 (4 out of 64 possible calls at each site), a minimum mean individual depth of coverage of 3 reads and a maximum of 80, and excluded all non-biallelic variants (1.7% of all variants). We used the program SnpEff (Cingolani *et al*. 2012) to annotate the variants and to predict the functional impact of each variant on protein coding genes and non-coding RNAs. High impact variants are defined as those predicted to cause premature start and stop codons, destruction of splice sites, and frameshift mutations in protein coding regions or non-coding RNAs.

In the high confidence variant set, we considered variants in difficult to sequence regions as missing data. Because *Arabidopsis* is highly inbreeding and expected to have high homozygosity throughout the genome, we identified genomic regions with low quality sequence by scanning for regions of unusually high heterozygosity indicative either of sequencing error or recent rare outcrossing events. Percent heterozygosity was calculated as the percentage of heterozygous genotype calls out of total calls at variant sites in 20 kb, non-overlapping windows. We calculated both within individual and across individual percent heterozygosity. In a single individual (line PA-DT1-12) we found large regions of high heterozygosity indicative of recent outcrossing, and considered genotype calls in these regions for this one individual as missing data. We also considered variants in windows with >20% heterozygous calls across all individuals as missing data. As predicted, heterozygosity was increased in highly repetitive centromeric regions of each chromosome and we excluded all variants in these regions (in total about 10.6 Mbp) from further analysis (equal to about 8.5% of the genome).

We used the program Pindel (Ye *et al*. 2009) to detect structural variants. We searched for insertions and deletions greater than 79 bp (the largest indel size detected in our genomes by the GATK). We also tested for reads supporting the presence of large replacements and tandem duplications. Structural variants were considered high confidence if they were supported in at least 4 individuals, and had a maximum mean depth of coverage below 80. We used a Fisher’s Exact Test to test whether each high confidence structural variant was over-differentiated between early and late flowering plant groups. Variants with p-values in the top 1% are shown in Figs S10 A-E.

### Population structure analysis

We used a subset of the high confidence SNPs to analyze population structure using the program Structure (Pritchard *et al*. 2000). We used only SNPs with a mean individual depth of coverage between 8 and 20 reads, and selected a subset spaced at least 10 kb apart throughout the genome to control for linkage disequilibrium (n=9852 SNPs). We used PGDSpider (Lischer and Excoffier 2012) to convert the SNP data to the correct format and then ran STRUCTURE using an admixture model with a burn in of 10,000 iterations, 10,000 sampling iterations, and K = 2 or 3 groups. The STRUCTURE model incorporated *a priori* information on sample population of origin.

To see how our sequenced lines were related to other accessions worldwide, we compared them to a subset of 90 geographically diverse lines genotyped by Platt *et al*. (2010). We used genotype information from whole genome sequencing calls at 135 SNP marker locations (Samis *et al*. 2012) to calculate distances between individuals and generate neighbor-joining trees in the program TASSEL (Bradbury *et al*. 2007). We included data for 65 lines scattered throughout Eurasia, 25 North American lines, and genotype calls from our 32 sequenced lines. To verify genetic diversity of our sample we also screened our samples for the presence of a common haplotype found in over 1000 lines from North America reported by Platt *et al*. [34]. That haplotype was represented in only 13% of our sequenced lines (n=4).

### Signatures of Selection

#### Differentiation as X^T^X (∼F_ST_)

We used four tests in genome-wide scans to detect regions of the genome with potential signatures of selection for flowering time. First, we identified variants that were over-differentiated between early and late flowering plants and also predicted to have deleterious impacts on protein coding genes and non-coding RNAs (as annotated by SnpEff, above). To measure differentiation we calculated the statistic *X*^*T*^*X* for high confidence variants using the program Bayenv2 (Günther and Coop 2013). The *X*^*T*^*X* statistic is analogous to F_ST_, but is based on allele frequencies that are normalized for population structure and uneven sampling between groups. The expected value of *X*^*T*^*X* is the number of groups being compared (n=2). We generated a kinship matrix used to normalize allele frequencies using a set of 3,000 high confidence synonymous coding SNPs spaced evenly throughout the genome. We then calculated *X*^*T*^*X* for two subsets of markers; 1) 20,213 high confidence synonymous SNPs spaced no closer than 2,500 bp from each other throughout the genome and 2) the set of all variants predicted to have strong deleterious impacts on genes (n=17,324). *X*^*T*^*X* values were calculated as the average value from 3 separate BayEnv2 runs using a burn-in of 30,000 iterations and 100,000 sampling iterations each. The distribution of values from synonymous SNPs was used to gauge background levels of differentiation between the early and late flowering plant groups. We considered SNPs over-differentiated if they had average *X*^*T*^*X* values above the average of the highest synonymous *X*^*T*^*X* values from across three runs. F_ST_ can be inflated by processes that reduce within-population genetic diversity (e.g., inbreeding depression, stronger background selection in some regions of the genome, differences in effective population size) that do not universally reduce variation in the entire sample (Charlesworth 1998). We do not expect this to be a substantial problem with our implementation of *X*^*T*^*X* because our comparison of early and late flowering genotypes incorporates samples from several populations, with different relationships to native samples, into the categories being compared. Consequently, the forces that would reduce within-sample genetic diversity in one flowering group but not the entire sample are unlikely to be occurring independently in several localities that make up the early or late-flowering groups.

#### Cross-population Composite Likelihood Ratio test

Second, we used the cross-population composite likelihood ratio (XP-CLR) test (Nielsen *et al*. 2005) to search for genomic regions indicative of selective sweeps. For this analysis we used the high confidence variant data set, but did not consider SNPs in high heterozygosity regions as missing data in order to avoid biases introduced by creation of large regions of missing genotype calls. We obtained physical and genetic map distances between 455 markers in the *Arabidopsis* genome from The *Arabidopsis* Information Resource (Rhee *et al*. 2003) (map files; RIdataGeneticMap.txt and TAIR9_AGI_Map_data). We used the genetic distance information from the markers to predict genetic distances between all markers in our populations using a 4^th^ order polynomial. We calculated both the XP-CLR of selection in the early flowering plant group using the late flowering group as a reference, and vice versa. Input genotypes were un-phased. The Loess smoothed prediction of these data is shown in Figs S10 A-E. The presence versus absence of a selective sweep was tested in 0.5 cM windows around grid points placed every 5kb along each chromosome. We excluded grid points within centromeric regions (due to poor genotyping accuracy in these repetitive areas). SNPs were down-sampled to a maximum of 100 SNPs per window. We calculated the difference between the XP-CLR statistic in the early group and the late group at each grid point, and considered as potential sweep regions those with difference values below the first quantile or above the 99^th^ quantile of values.

#### Diversity statistics: Π and Tajima’s D

Lastly, we looked genome-wide for regions of unusually low nucleotide diversity (π) and low values of Tajima’s D (Tajima 1989) in either the early or late flowering plants. Measures of nucleotide diversity are sensitive to the accuracy of genotype calling at invariant sites, so we used depth of coverage information at invariant sites as a measure of genotype quality in the analysis. To generate information for invariant sites throughout the genome we used a similar but alternative GATK pipeline to the one previously described to call variants. Alignments were used to call variants separately for each individual against the reference genome using GATK Haplotype Caller. A combined variant set was then generated using GATK CombineGVCFs followed by GenotypeGVCFs, emitting genotype and read depth information for every base in the genome.

We calculated π and Tajima’s D in fixed windows across the genome using only genotype calls with a minimum depth of 4 reads. To keep the number of individuals sampled at each site constant, we randomly subsampled genotype data for 12 individuals only at sites where at least 12 individuals had non-missing calls. π and Tajima’s D were calculated for windows containing 10,000 bases meeting the read depth and missing-data cutoffs within each phenotypic group. Using the window data, we found the difference between π values in the early and late flowering lines at each grid point previously used to calculate XP-CLRs. The same was done for Tajima’s D. We considered grid points with difference values below the first quantile and above the 99^th^ quantile of all difference values as potential sweep regions in the late and early flowering groups, respectively.

### Identification of genes potentially under selection

We examined a list of 284 candidate genes, identified by Brachi *et al*. (2010) and Atwell *et al*. (2010), which have annotations broadly associated with plant development and developmental responses to endogenous and exogenous cues. We also included in the list three genes identified as associated with flowering time by Atwell *et al*. (2012). Genes in this list we term ‘*a priori*’ candidate genes for flowering time. For each candidate locus, within each phenotypic group, we calculated π and Tajima’s D using non-missing genotype calls with data for at least 7 individuals. When more individuals had valid calls the data was randomly down sampled to 7 within each phenotypic group. We then compared π and D values for each gene between the early and late flowering plant groups, and compared the differences against the distribution of differences calculated similarly for all other genes in the genome.

To detect additional genes potentially under selection for flowering time, we identified genes that fell in regions of the genome characterized by signatures of selection described above (π, Tajima’s D, XP-CLR). Regions of interest were defined using the window boundaries closest to the significant gridpoints, and all genes overlapping those regions were identified. XP-CLR regions of interest were defined as 2500 bp up-stream and down-stream of significant gridpoints. The strongest non *a priori* candidate genes for involvement in flowering time adaptation were in genomic regions with more than one signature of selection or had both a signature of selection and contained over-differentiated Fst (*X*^*T*^*X*) outliers. We also conducted functional term enrichment analyses on genes with one or more signature of selection using the online tools David (Huang *et al*. 2009) and GOrilla (Eden *et al*. 2009). The set of all genes affected by over-differentiated (but non-deleterious) SNPs was used as the background set for enrichment analyses of high impact outliers. The set of all *Arabidopsis* genes was used as the background set for enrichment comparisons of genes in regions with significant signatures of selection.

### Extended analysis of candidate loci FLC, FRI, PHYC, and MAF2

Indel and SNP polymorphisms in the loci FLC, FRI, PHYC, and MAF2 have been previously linked to geographic variation in flowering time in *Arabidopsis* (Caicedo *et al*. 2004, 2009; Balasubramanian *et al*. 2006; Samis *et al*. 2008; Rosloski *et al*. 2010; Li *et al*. 2014). We compared the sequences of these three loci and the surrounding regions with sequences published in other studies to determine if 1) polymorphisms associated with flowering time identified in other studies are present and similarly associated in the 32 North American lines, and 2) if the genes or surrounding sequence haplotypes at these loci show any structuring between early and late flowering lines. We extracted the sequence of each locus from each individual, counting bases without any read coverage as missing data. For +/- 50kb regions surrounding each locus, polymorphisms were extracted and concatenated. Genic sequences were used, as aligned to the reference genome, to create neighbor-joining trees (model = “raw”) using the ‘ape’ package in R, with pairwise.deletion = TRUE (Paradis *et al*. 2014).

### Data Availability

*Arabidopsis* lines are available on request. Table S2 contains geographic origin information on the sequenced lines. Short-read sequence data for each line will be deposited in the NCBI short-read archive (SRA) under BioProject number PRJNA288374. A VCF file containing genotype information on the sequenced lines is available on request.

## Results

### Unique genetic diversity and evidence of outcrossing in North American populations

We found introduced *Arabidopsis* populations have substantial genetic diversity, much of which is unique to the introduced range or at least uncommon in Eurasia. We compared the approximately 2.35 million variants in our initial data set (those with non-missing genotype calls in at least 50% of individuals) with available large polymorphism data sets for *Arabidopsis* including data from the 18 Eurasian parental accessions used to generate the MAGIC nested association reference panel (Gan *et al*. 2011), the first 80 genomes from the 1001 genomes project (Cao *et al*. 2011), and ∼215,000 variants generated from the worldwide RegMap panel (Horton *et al*. 2012) (Fig. 1). Twenty-seven percent of the variable sites in our North American lines (n=∼644,000) were only present in our data set and not in any other. The RegMap variant data, one of the few available genomic data sets containing a significant fraction of North American lines, captured only 906 (0.1%) of the variants unique to the species introduced range.

**Figure 1.**
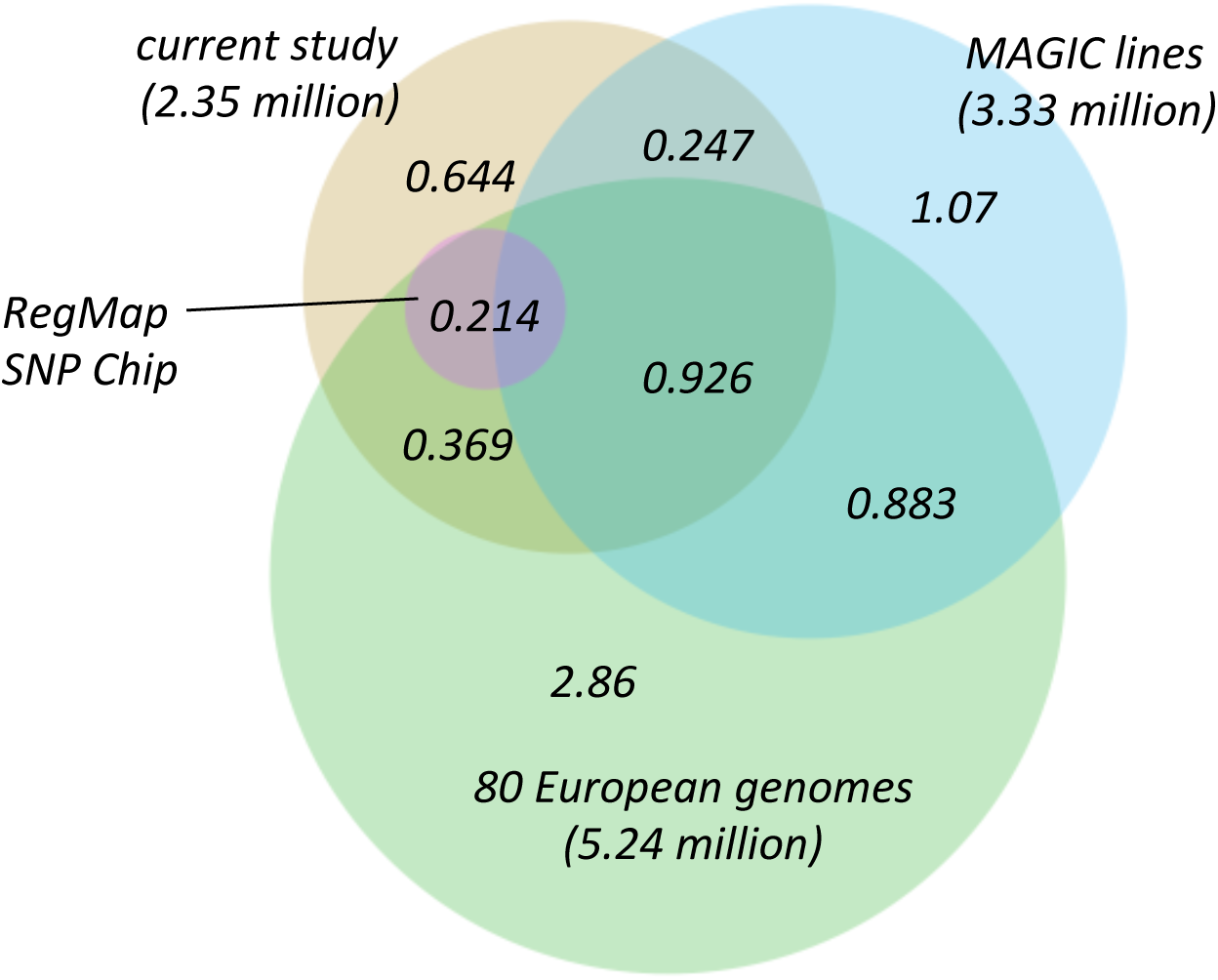
Overlap between variants identified in the current study and 3 other studies. (Cao *et al.* 2011; Gan *et al.* 2011; Horton *et al.* 2012).

We used a subset of the highest confidence SNPs to estimate population genetic parameters for eastern North America. On average, short-range LD dropped off at approximately 7 kb on each autosome which is similar to LD estimates of 10 kb in the native range (Kim *et al*. 2007). Based on average pairwise diversity of our sample, the effective population size (N_e_) for eastern North America is about 215,000 which is comparable with regional estimates of N_e_ in Europe (Cao *et al*. 2011). Fixed-window analysis of individual genomes revealed wide regions of unusually high heterozygosity in a single individual from Pennsylvania (PA:DT1-12, Fig. S2) on chromosomes 1, 4, and 5, which is indicative of a recent rare outcrossing event.

### Population structure consistent with multiple introductions and mixing among early and late flowering groups

Genome-wide data on ∼9,800 SNPs showed that natural accessions in the introduced range cluster genetically by geographic sub-region, with some exceptions (Fig. 2A and S3). Some individuals did not cluster with their nearest geographic neighbors and others were likely to be admixed. For example, two lines from a single population in Massachusetts (HVSQ) each clustered strongly with a different ancestral genetic group (Fig S3). Some lines from the Midwest (populations Knox and Kin) and from New York (population CSH) also had mixed ancestry.

**Figure 2.**
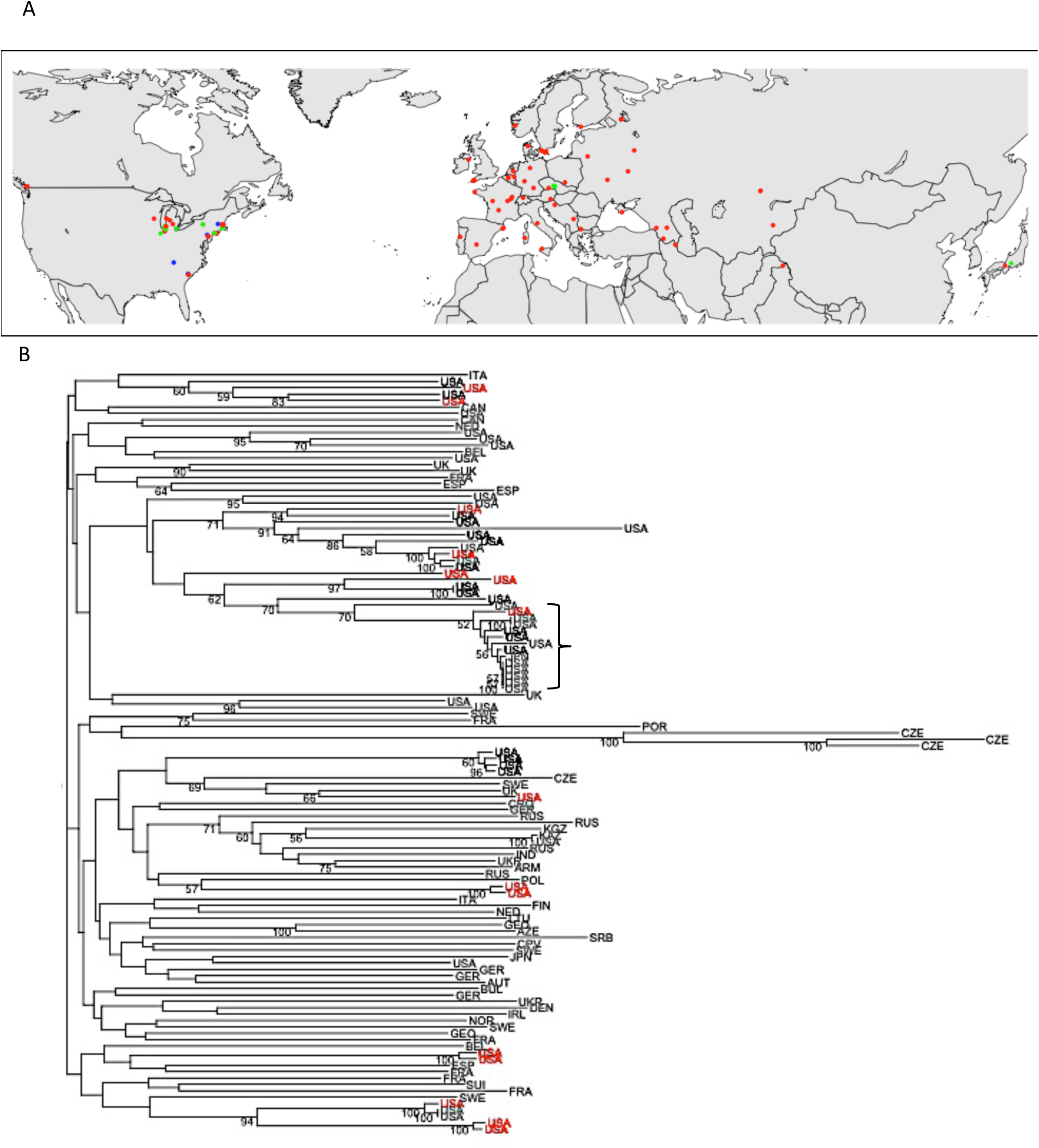
Popula5on structure and loca5on of sequenced lines. A) The color of each marker represents the Structure group to which each line most likely belongs (K=3 groups). **B)** Neighbor-joining tree based on 126 SNP markers. Early flowering lines are in red, late flowering lines are in black, lines genotyped by PlaQ *et al.* [34] are in gray. Lines are labeled with their country of origin. The bracket shows individuals with the most common North American haplotype detected by [34]. Bootstrap percentages are shown for nodes >50%.

Comparison of North American lines and a subset of natural accessions from around the world (compared at 126 SNP markers genotyped by Platt *et al*. [34]) showed North American populations cluster with lines from many different parts of Eurasia (Fig 2). Clustering was consistent with previous analyses of smaller SNP sets which indicate patterns of isolation by distance at the scale of ∼250 km and at least two ancestral lineages in North America (Kim *et al*. 2007; Beck *et al*. 2008; Platt *et al*. 2010; Samis *et al*. 2012). The analysis of additional genome-wide polymorphisms indicated the likelihood of only one additional ancestral group (K=3) (Fig. S3). There was no indication that early and late flowering genotypes stem from independent lineages (Fig S1 B).

### Outlier analysis reveals deleterious variants associated with flowering time

We determined whether any of the high confidence variants with predicted deleterious impacts on protein coding genes were over-differentiated between the early and late flowering plant groups and thus potentially causative of phenotypic differences. We considered the impact of a variant potentially deleterious to the protein product of a gene if it caused a frame-shift mutation, a gain or loss of a stop or start codon, or disrupted splice sites. As a measure of differentiation between phenotypic groups at each deleterious variant we used *X*^*T*^*X* (an F_ST_-like statistic) implemented in the program BayEnv2 (Günther and Coop 2013). *X*^*T*^*X* controls for residual population structure through the use of a kinship matrix generated from a subsample of genome-wide SNPs. In our simulations, the mode of the distribution of *X*^*T*^*X* values for unlinked synonymous coding SNPs throughout the genome was 1.96, close to the expected value for neutral variants (*X*^*T*^*X* =2.0), demonstrating adequate control for population structure. In contrast, SNPs predicted to have deleterious impacts on proteins had a right skewed distribution with modal *XTX* = 2.8 (Fig S4). The maximum *XTX* value for synonymous SNPs was 2.8 but high impact variants had values up to 8.1, indicating some are likely to be under selection. There were 996 deleterious variants with differentiation values above the maximum observed at any synonymous SNP and these had putative impacts on 416 protein coding genes. The set of impacted genes was not significantly enriched for any gene ontology terms related to physiology or environment. Among a list of 284 genes with functions related to flowering time, development, or environmental sensing (termed ‘*a priori*’ flowering time genes, see Methods), we looked to see if any contained over-differentiated deleterious SNPs. Five *a priori* flowering time candidates contained such variants, however this number is not more than expected by chance (χ^2^=1.647, p = 0.10). Among the genes containing the most differentiated non-synonymous SNPs (n=156) between the two phenotypic groups there was also no significant enrichment for *a priori* flowering time candidate genes (p=1.0) or GO terms related to development or environmental sensing.

We also scanned the genome for large structural variants and tested whether any were similarly associated with flowering phenotype. We used the program Pindel to search for variants larger than the largest insertion or deletion detected using the GATK variant calling pipeline (72 bp). There was strong support for the presence of 3,062 large structural variants in the genome, almost 75% of which were deletions with respect to the reference (Fig. S5). Forty-three structural variants were highly differentiated between early and late flowering genotypes (Fisher’s Exact test, lower 1% quantile of p-values, p<0.0002) and these were within 5 kb of 60 genes. One affected gene was an *a priori* flowering time candidate gene, the floral organ identity gene Enhancer of HUA2 (HEN2).

### Targeted analysis of flowering time candidate genes shows little evidence of selection or association with flowering variation

We next identified loci with the largest difference in nucleotide diversity (π) and Tajima’s D between the early and late flowering plant groups. A reduction in diversity or Tajima’s D in one group relative to the other may indicate recent selection on that locus in the group where the statistic is lowest. Only one out of 284 *a priori* flowering time candidate genes fell within the top 1% of genes with reduced diversity (π) in late flowering lines, a subunit of the transcription factor nuclear *factor Y* (NF-YA3). Similarly, a single *a priori* candidate gene fell within the top 1% of genes with reduced diversity in early flowering lines, the transcription factor tiny (TNY) (Fig. S6 A). For Tajima’s D, only two candidate loci showed evidence of statistical differences between the early and late flowering groups beyond the 1% threshold, the floral repressor *schlafmutze* (SMZ) and *phosphoglucomutase* 1 (PGM) (Fig S6 B).

In the native range of *Arabidopsis* at least four genes have alleles that affect flowering time and also occur at moderately high frequencies across the range: FLC, FRI, PHYC, and MAF2. We evaluated whether the known common variants at these loci are present in North America and associated with flowering time differences. We also tested for haplotype structuring at these loci indicative that nearby variation is associated with phenotype. We did this by constructing phylogenetic neighbor-joining trees based on alignments of the coding regions and regulatory regions (50 kb up- and down-stream) of these genes (Fig S7).

Full sequences of the FRI and FLC genic regions confirmed the results of previous PCR-based genotyping (Samis *et al*. 2012). FRI contains a ∼376 bp deletion that was present in 8 lines, and a 16 bp insertion present in all but 2 lines. Neither deletion was strongly associated with flowering time phenotype and analysis of structural variation using Pindel (Ye *et al*. 2009) did not detect any additional large variants in FRI. We observed several other small putative deletions affecting FRI, but all occurred at low frequency (in 1 or 2 individuals).

Within FLC, we did not detect any of the large transposable element insertions that occur within the FLC^A^ group (or any other large structural variants within the gene) and confer early flowering in the Eurasian range (Gazzani *et al*. 2003; Michaels *et al*. 2003). Haplotypes in the 100 kb region surrounding FLC did not closely match any of the European rapid or slow vernalization haplotypes detected by Li *et al*. (Li *et al*. 2014) with the exception of a single early flowering North American line that had strong similarity to the early flowering rapid-vernalization-2 haplotype (RV2, Fig S8).

The photoreceptor phytochrome C (PHYC) contains alleles that vary longitudinally in frequency across Europe and affect flowering in response to day length in some mapping populations (Balasubramanian *et al*. 2006; Samis *et al*. 2008). PHYC is characterized by Ler-type and Col-type haplotypes that are distinguished by a large deletion associated with early flowering and sensitivity to day length. Analysis of PHYC haplotypes in North America showed that lines can also be distinguished as Ler- and Col-types, although neither haplotype is strongly associated with early or late flowering (Fig S9). Four lines from Rhode Island form a unique group outside of the Ler- and Col-PHYC types, and all are late flowering, however the association may be due to population structure rather than the presence of a late flowering functional variant near PHYC. The 100kb region surrounding PHYC is highly polymorphic containing 7 or more haplotype clusters, none of which are characterized by a single flowering time phenotype (BG and JRS, unpub. data).

Lastly, we examined variation in *MADS AFFECTING FLOWERING 2* (MAF2). Large insertions in MAF2 originating from an adjacent tandemly duplicated paralog, MAF3, are common throughout Eurasia and associated with delayed flowering (Caicedo *et al*. 2009; Rosloski *et al*. 2010). Sequence alignments showed no evidence of MAF2 insertions in any of the North American lines outside of two regions of microsatellite variation. Neither genic nor upstream MAF2 haplotypes clustered by flowering time phenotype, and Pindel analysis did not detect any large structural variation within the locus.

### Intersection of selection scan results identifies genes in low diversity genomic regions

We calculated nucleotide diversity (π), Tajima’s D (TAJIMA 1989), and the cross population composite likelihood ratio (XP-CLR) in non-overlapping windows across the genome to detect regions potentially affected by recent sweeps due to selection for flowering time. The presence of a recent sweep is indicated where there is a large difference in nucleotide diversity, Tajima’s D, or the XP-CLR between the early and late flowering groups relative to the distribution of difference values in other regions throughout the rest of the genome (Fig S10). The top 1% of genome-wide windows where signatures of selection had the greatest difference between early and late flowering lines, contained 1884 genes (Fig. 3). These genes are potential candidates for involvement in flowering time evolution, but the set was not enriched for *a priori* flowering time candidates (n=27, Fisher’s Exact test, p=0.10). There was also no significant enrichment for gene-ontology (GO) terms related to development or environmental sensing.

**Figure 3.**
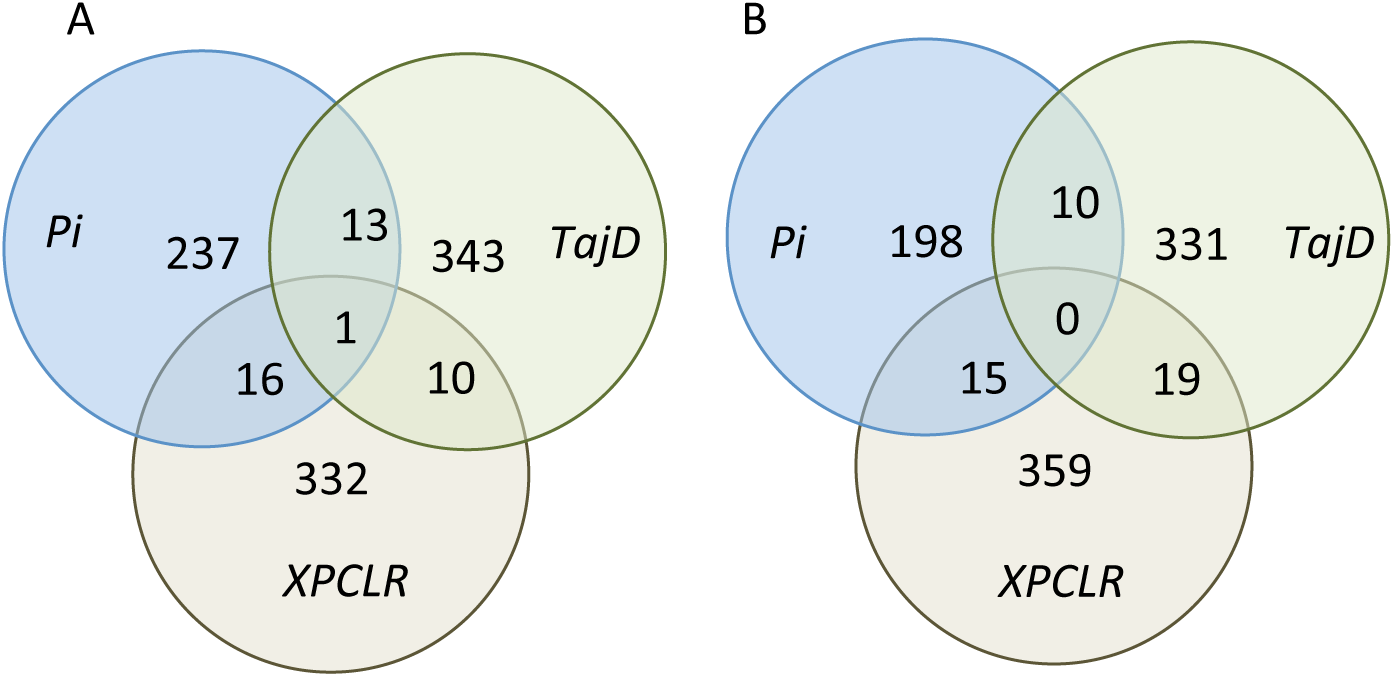
Genes in genomic regions with signatures of selec5on. A) selection for early flowering; B) selection for late flowering.

Of the 1884 genes that fell in regions of the genome indicative of potential selection on flowering time, 43 contained one or more variants predicted to have deleterious impacts on protein function and that were also over-differentiated between early and late flowering plants (*X*^*T*^*X* > 3.48; Fig. 4, Table S1). Ten additional genes from the set of 1884 were in close proximity to over-differentiated structural variants (Table S1). These 53 genes are candidates for involvement in flowering time differentiation in the introduced range.

**Figure 4.**
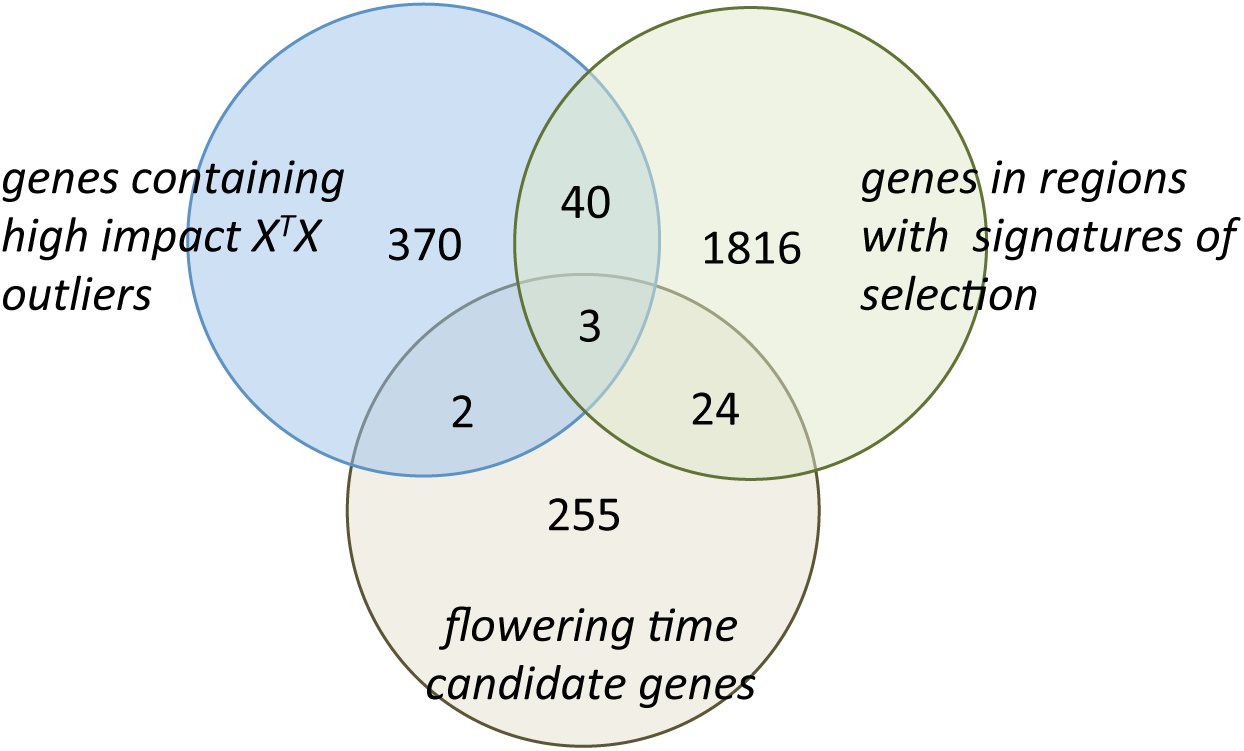
Genes with evidence of involvement in flowering 5me evolu5on. The overlap is shown between genes containing highly differentiated SNPs with predicted deleterious impacts (*X*^*T*^*X* outliers), genes in regions with a signature of selection for flowering time (Fig. 3), and genes identied as *a priori* candidates based on functional annotation (from [23]).

For the 43 candidate genes with over-differentiated, deleterious SNPS, we measured the frequency of the alternate allele in both our sample and in Europe (using data generated by Cao *et al*. (30)). Most of the deleterious indels had zero frequency in the European data set, but this is most likely due to low repeatability between current indel prediction methods. A more reliable comparison between allele frequencies can be made between the over-differentiated deleterious SNPs (n=18) in the two ranges. Among these, the frequency in Europe was predictive of the frequency in North America (slope = 0.72, p=0.0008, R^2^=0.49, Fig. S11). However, the SNPs were not more or less common in the native than the introduced range.

## Discussion

Whole genome sequencing in natural *Arabidopsis* accessions from eastern North America has revealed little evidence that the evolution of flowering time differences in the introduced range has involved selection on known genetic variants from the native range. Some mutations known to have strong influence on flowering time do occur in the introduced range, but we found no evidence that these mutations are associated with differences in flowering time in introduced populations or have been subject to recent hard selective sweeps. Further, regions of the genome containing the strongest signatures of selection in early or late flowering lines do not contain genes that are enriched for functions in development or sensing of environmental cues. Thus, it is unlikely that the evolution of flowering time in the two parts of the range is strongly parallel at the level of genes. Our results, therefore suggest, that neither pleiotropic constraint nor available standing genetic variation has constrained the evolution of flowering time to selection on high frequency functional mutations from the native range or known flowering time pathway genes.

We identified regions of the genome with large differences in diversity (π) or Tajima’s D between phenotypic groups, however on average differences in the top 1% of regions were small and consistent with only putatively weak selection. For signatures calculated for individual genes, few *a priori* flowering time candidates fell within the top 1% of values. Given the diversity of introduced populations and evidence of multiple introductions from this and other studies, standing genetic variation has most likely played a central role in the evolution of traits in the introduced range. The absence of a signal of strong selective sweeps at flowering time genes in the introduced range thus might be explained by several non-mutually exclusive mechanisms: 1) evolution through soft rather than hard sweeps at candidate or other loci; 2) polygenic adaptation involving many genes of small effect; and 3) selection on rare alleles present at low frequency in our sample.

### Possible soft selection for flowering time

Soft selection on standing genetic variation is probable in introduced *Arabidopsis* which has a history of multiple introductions from different parts of Eurasia and thus probable introduction of multiple adaptive alleles at many loci. Signatures of selection on genes with multiple adaptive alleles may not be strong enough to detect with diversity-scans. While diversity and site frequency spectrum deviations around loci subject to soft selective sweeps may not be differentiable from drift, haplotype structuring around such loci may reveal phylogentic grouping by phenotype (Messer and Petrov 2013). Statistical analysis can reveal such patterns genome-wide (Sabeti *et al*. 2007) as can tree-based analyses of specific candidate loci. Haplotype structuring around candidate genes FRI, FLC, PHYC, and MAF2 did not reveal any evidence of haplotype clustering by phenotype in the introduced range, however some structuring was apparent at two developmental transcription factor genes found in regions of the genome with the strongest selection signatures (TFIIIA and NF-YA3, Fig S12). Soft sweeps at these and other loci may have contributed to flowering time in the introduced range and warrant further investigation.

### Polygenic adaptation

Likewise, allele frequency changes at many genes of small effect on flowering time may go undetected in outlier scans. The importance of polygenic adaptation in evolution in response to new environments is currently unknown, but some have argued may be the predominant form of quantitative trait evolution from standing genetic variation (Hancock *et al*. 2010; Pritchard and Di Rienzo 2010; Pritchard *et al*. 2010). We found no positive evidence of polygenic flowering time adaptation in introduced populations in the form of enrichment analyses. Enrichment analyses provide a way to search for a signal that many loci have facilitated adaptation of flowering time. No set of genes putatively under selection identified in this study was enriched for GO terms related to development or environmental sensing. This includes genes containing the most differentiated non-synonymous snps, genes containing high-impact *X*^*T*^*X* outliers, and genes in regions with putative signatures of selection. However, it is highly likely that our study lacked statistical power to detect subtle allele frequency changes across many genes characteristic of polygenic adaptation. For example, in studies of height in humans, (Lango Allen *et al*. 2010; Turchin *et al*. 2012), the allele frequency differences observed are often quite small. For instance, in [44] allele frequency differences between Northern and Southern human populations ranged from 0.0044 to 0.016 with sample sizes ranging from 58 to 257 in each population. Consequently, while a significant enrichment test could have been interpreted as evidence in favor of polygenic adaptation, its absence cannot be used as evidence against it given sample sizes.

In addition to power considerations, polygenic adaptation may contribute in ways that are difficult to discern from genomic portraits of polymorphism and divergence. Theoretical work by Latta (Latta 1998, 2004) and LeCorre and colleagues (Kremer and Le Corre 2012; Le Corre and Kremer 2012) has demonstrated that trait divergence can evolve even with limited differentiation of the underlying causal loci. In brief, because selection generates covariances among different loci, changes in genetic variances, and hence quantitative genetic differentiation, can far outpace allele frequency changes in individual loci. Consequently, outlier-based approaches, such as ours, are likely to represent a conservative lower bound of the loci involved in adaptation.

### Selection on locally common but regionally rare alleles

In the absence of any strong evidence for a parallel genetic basis of flowering time clines, selection on locally common but regionally rare alleles may explain a portion of the evolution of flowering time in the introduced range. Adaptive alleles and haplotypes that occur only in small areas of the introduced range would not occur often enough within our set of introduced lines to show a signal of over-differentiation or reduced diversity. In the native range, previous studies have shown that alleles that are regionally restricted in their distribution explain up to 24% of the variation in some fitness related traits (Fournier-Level *et al*. 2011; LI *et al*. 2014). The alleles are not associated with signatures of selection, are rarely associated with fitness in more than one region, and their distribution is also often spatially correlated with climate factors. In the introduced range, the evolution of flowering time may follow a similar pattern and would be facilitated by the introduction of adaptive alleles from throughout the native range. For example, we found some evidence of uncommon deletions in Fri in our sample, some of which could be at high frequency at a local scale and contribute to flowering time phenotype only in certain populations. In a similar case, various different deletions in a cyanogenesis gene have recently been found to contribute to the development of a cyanogenesis cline in wild clover (Kooyers and Olsen 2012; Olsen *et al*. 2013). In the future, greater within-population sampling of *Arabidopsis* in North America will be able to test the hypothesis that adaptive alleles are regionally rare but locally abundant.

### New candidates for flowering time evolution in the introduced range

Despite the limitations of outlier-based genome scans, many of the loci detected in the present study are likely to represent true recent targets of selection. Outlier-based selection scans are most powerful at detecting selection that results from newly arising mutations or similarly, those that exist within bottlenecked populations prior to demographic expansion (Teshima *et al*. 2006; Jensen *et al*. 2007; Pritchard and Di Rienzo 2010; Lotterhos and Whitlock 2015), and the signature of full or partial selective sweeps in the genome are expected to persist on the order of N_e_ generations (Przeworski 2002). Introduced *Arabidopsis* has a relatively large effective population size in eastern North America (comparable to regional diversity in Europe, N_e_ ∼215,000 in this study), and a history of very recent introduction over the past ∼250 years (Jørgensen and Mauricio 2004). Prior to population expansion, sweeps should be detectable but rare in the introduced range. Results of outlier tests are not likely to detect all targets of selection for flowering time evolution in the introduced range, but the use of multiple test statistics, as in this study, encourages a high rate of true positives among outlier loci [53]. The genes that fall within regions of the genome with the strongest signature of flowering time selection and that also contain over-differentiated, high-impact deleterious variants are the best candidates for involvement in flowering time evolution in the introduced range. This is a set of 43 genes impacted by over-differentiated variants (Table S1), and includes 3 *a priori* candidate genes: NF-YA, TFIIIA (mentioned above), and TNY (a DREB transcription factor). It also includes two heat-shock proteins, a thigmomorphogenesis locus (involved in response to mechanical stimuli), and a MADS-box transcription factor. We also consider as candidate genes the 10 genes in close proximity (5kb) to large over-differentiated structural variants that also lie in regions with a strong signature of selection for flowering time (Table S1). In both sets of candidate genes, some loci have putative developmental functionality, however none have alleles of which we are aware that are known to influence flowering time in the species native range.

Non-parallelism at the level of genes in the evolution of flowering time clines in the native and introduced range of *Arabidopsis* is consistent with the results of a recent study by Hamilton and colleagues (2015) (Hamilton *et al*. 2015). In a transplant of Eurasian genotypes to a common garden in Rhode Island, they found that adaptive variants in the native range differed from adaptive variants in Rhode Island, despite similar climate conditions. Most of the adaptive alleles in Europe and Rhode Island exhibited conditional neutrality: beneficial in one site, with little or no effect in the other. The prevalence of conditional neutrality underlying local adaptation has important consequences for evolution in introduced species. If loci underlying local adaptation in the native range frequently exhibit conditional neutrality (Anderson *et al*. 2011, 2013; e.g. Fournier-Level *et al*. 2011) one would predict that they would be locally abundant, but possibly not have spread range-wide. For species with a history of multiple introductions, like *Arabidopsis*, a complex mosaic of previously locally adapted yet globally rare alleles may underlie any post-introduction evolutionary responses. In these scenarios, evolution of similar broad geographic patterns through parallel genetic mechanisms may be unlikely.

Collectively, our results show that post-introduction evolutionary responses are neither confined by pleiotropic constraint to a handful of loci previously implicated in flowering time evolution, nor constrained by limited standing genetic variation. Further studies on the genetic basis of adaptation in complex, quantitative traits will be necessary to determine the prevalence of this pattern.

## Acknowledgements

We would like to thank Adriana Salcedo, Stephen Wright, Wei Wang, Robert Williams, Emily Josephs, and Maggie Bartkowska for help and advice with analyses. This project was funded by NSERC Canada.

